# LFPy – multimodal modeling of extracellular neuronal recordings in Python

**DOI:** 10.1101/620286

**Authors:** Espen Hagen, Solveig Næss, Torbjørn V. Ness, Gaute T. Einevoll

## Abstract

LFPy is an open-source tool for calculating brain signals such as extracellular potentials and magnetic signals from simulated activity in multicompartment neuron models, ranging from single cells to large neuronal networks. The tool is provided as a Python package, and relies on the NEURON simulation environment.

## 2. Introduction

Electric extracellular measurements of neuronal activity in brain tissue has been one of the main workhorses in experimental neuroscience for several decades. Although such measurements are relatively easy to carry out, the large number of contributing sources to the measured signals renders the interpretation of the experimental data difficult. On the macroscopic scale, such as for measurements of the electric potential on the scalp, that is, electroencephalography (EEG) (Nunez and Srinivasan, 2006), or measurements of magnetic fields outside the head itself, that is, magnetoencephalography (MEG) (Hämäläinen et al., 1993), the interpretation is further complicated by the fact that the measurements are performed some distance from the neural tissue itself. EEG signals are also affected by the different electric conductivities of tissue between the brain and the electrodes (cerebral spinal fluid – CSF, bone and soft tissues) (Nunez and Srinivasan, 2006).

The availability of biophysics-based *forward*-modeling tools is important for hypothesis testing with precisely defined mathematical models, where the experimental system, including neurons and measurement devices, are mimicked in the virtual model and simulated on a computer. Construction and improvements to *inverse* analysis methods applied to experimental data (that is, inferring underlying neural activity from measurements) also calls for improved forward-modeling tools. Inverse methods are required to better understand and analyze the link between measured brain signals and the underlying neural activity. Forward-modeling tools allows for models of measurements with known underlying neural activity, ‘ground truth’, that inverse methods can be validated against.

LFPy (LFPy.readthedocs.io) implements different established forward models for extracellularly recorded potentials and magnetic fields, stemming from activity in multicompartment neuron models (De Schutter and Van Geit, 2009). Extracellular potentials in vicinity of the neurons are computed as distance-weighted sums of contributions from the transmembrane currents of the simulated neurons, while magnetic fields are either calculated from axial currents within the neurons or via the so-called current dipole moment calculated from the associated transmembrane currents. The current dipole moment is also used for EEG predictions. The corresponding electrostatic and magnetostatic forward models are derived using the appropriate volume-conductor theory (Hämäläinen et al., 1993; Nunez and Srinivasan, 2006; Einevoll et al., 2013).

## 3. Application

The first version of LFPy (Lindén et al., 2014, v1.0) incorporated the well-established scheme for extracellular potentials pioneered by Holt and Koch (1999), where transmembrane currents of multicompartment models are first computed using a domain-specific software (NEURON Simulation Environment, Carnevale and Hines (2006)). Then the extracellular potential is computed as the distance-weighted sum over each current contribution, with weights derived using theory for an infinite homogeneous, isotropic and ohmic volume conductor model.

This numerical scheme has been applied to predictions of the local field potential (LFP), that is, the low-frequency part of the extracellular potential (Pettersen et al., 2008; Lindén et al., 2010, 2011; Gratiy et al., 2011; Makarova et al., 2011; Schomburg et al., 2012; Łęski et al., 2013; Reimann et al., 2013; Martín-Vázquez et al., 2013, 2015; Gląbska et al., 2014; Mazzoni et al., 2015; Tomsett et al., 2015; Sinha and Narayanan, 2015; Taxidis et al., 2015; Hagen et al., 2016; Gląbska et al., 2016; Ness et al., 2016, 2018; Hagen et al., 2017; McColgan et al., 2017), as well as extracellular spike waveforms which carry more power at high frequencies (Holt and Koch, 1999; Gold et al., 2006, 2007; Pettersen and Einevoll, 2008; Pettersen et al., 2008; Franke et al., 2010; Schomburg et al., 2012; Thorbergsson et al., 2012; Reimann et al., 2013; Ness et al., 2015; Hagen et al., 2015; Miceli et al., 2017; Cserpán et al., 2017; Luo et al., 2018; Buccino et al., 2018, 2019).

Since LFPy 2.0 (Hagen et al., 2018), the tool has been extended to also support forward-model predictions accounting for anisotropic media, that is, with different conductivity in different directions (Goto et al., 2010), and discontinuous media where the conductivity is piecewise constant in a single direction. The latter can for example be used to mimic in vitro experimental setups using microelectrode arrays (MEAs) (Ness et al., 2015), or to mimic in vivo conditions where a jump in conductivity can be expected such as at the boundary between the brain and CSF. Furthermore, LFPy now incorporates calculations of axial currents across the neuronal morphology for estimating the magnetic field nearby the neuron (Blagoev et al., 2007). Hence, corresponding experiments using magnetic detection devices directly in neural tissue (Barbieri et al., 2016; Caruso et al., 2017) can be mimicked.

LFPy also features calculations of current dipole moments from transmembrane currents (Lindén et al., 2010), which is used for calculation of both EEG and MEG signals. For EEG predictions LFPy incorporates the analytical 4-sphere head model (Nunez and Srinivasan, 2006), which was corrected by Næss et al. (2017), to account for the different conductivities of brain, CSF, skull and scalp. The computed current dipole moments can also be used with more complex head models (see e.g., Huang et al. (2016)). MEG signal predictions in a spherically-symmetric head model rely on a special form of the magnetostatic Biot-Svart law (ignoring effects of magnetic induction) (Nunez and Srinivasan, 2006, Appendix C).

Finally, in contrast to its first release where the tool only supported simulations with a single cell instantiation at the time, LFPy also supports networks of synaptically interconnected neurons. Such network simulations can be executed on a single physical machine, but larger network simulations is better executed in parallel on high-performance computing (HPC) facilities (Hagen et al., 2018).

## 4. Architecture

LFPy is a Python package that presently (Hagen et al., 2018) provides different high-level class definitions which represent cells, populations and networks of cells, synapses, intracellular stimulation devices (current and voltage clamps) and extracellular recording devices (representing both invasive and noninvasive equipment). LFPy’s class definitions rely internally on NEURON’s Python interface (Hines et al., 2009). As many illustrative LFPy usage examples are provided online (github.com/LFPy/LFPy/examples), we here only briefly summarize the main classes and their intended use, and provide a simple use case below. The best way to learn the tool is by going through the various example files. More detailed technical documentation is available online^1^.

The most basic neuron representation in LFPy is provided by the `LFPy.Cell` class. A `Cell` object is typically instantiated with a chosen morphology, either on the form of a morphology file with instructions NEURON can digest (with file endings ‘.hoc’, ‘.asc’, ‘.swc’ or ‘.xml’), or a NEURON `SectionList` instance filled with `Section` references. A number of optional arguments can be used to set the passive parameters of the cell model, additional routines to distribute for example active ion channels to different parts of the neuron, rules for spatial discretization of the corresponding cable model, and simulation settings (temporal duration and discretization). After instantiation, the cell object has several public methods that can be used to position and align the cell in space, return indices of different compartments of the model, and run the simulation.

Two additional classes are defined using inheritance from class `Cell`, named `TemplateCell` and `NetworkCell`. These can be used with existing ‘network-ready’ single-cell models that uses NEURON’s *template* definitions which allows for multiple concurrent instantiations of individual neurons. In LFPy however, the `TemplateCell`’s intended use is still for handling single-neuron simulations (serially), while `NetworkCell` objects are instantiated in neuronal network populations.

After a cell’s instantiation, it can be instrumented with synapses by the creation of an arbitrary number of instances of class `LFPy.Synapse`. The main parameters of the `synapse` class is a reference to the cell object itself, the compartment index where the synapse is located, and synapse type with synapse-specific parameters (time constants, weight, reversal potential etc.). The synapse current and postsynaptic potential can optionally be recorded during the course of a simulation. The instantiated synapse has public methods to set its activation times, to either explicit values or determined stochastically using NEURON’s `NetStim` class.

Intracellular stimulation devices (current and voltage clamps) can be set up using LFPy’s `StimIntElectrode` class. The class is instantiated similar to synapses, but the name of point-process type has to be defined as well as its specific parameters. Recording of stimulation current and corresponding compartment membrane voltage can optionally be enabled.

The creation of networks, or simply different populations of concurrent `NetworkCell` instantiations, is in LFPy incorporated through its `Network` class. Its main parameters are the temporal discretization and duration of the simulation, the globally shared initial voltage for compartments and simulation temperature which affects the active ion-channel dynamics. The network instance contains a public method to create populations of the same cell type (through class `Population`), that is, with the same parameters being passed to the `NetworkCell` instances upon their creation. The network also contains a public method to connect pre-and postsynaptic populations using shared synaptic parameters. Values for connection weights, delays and synaptic locations can however be drawn randomly from desired function declarations such as `np.random.normal` with corresponding parameters, or be set to constant values.

### 4.1 Example code

Here, we demonstrate how to set up a simple LFPy simulation in Python. We define a simplified neuron representation with an active soma compartment and passive apical dendrite which receives strong synaptic input at the terminating end of the dendrite, with simultaneous calculation of the extracellular potential alongside the neuron. We first create a ‘ball and stick’ morphology file, with a passive dendrite and active soma using NEURON’s HOC syntax from Python:

~~~
morphology = ‘BallAndStick.hoc’
with open(morphology, ‘w’) as f:
 f.writelines(‘‘‘// Create sections:
create soma[1]
create apic[1]
// Add 3D information:
soma[0] {
 pt3dadd(0, 0, −15, 30) // x,y,z,d in um
 pt3dadd(0, 0, 15, 30)
}
apic[0] {
 pt3dadd(0, 0, 15, 3)
 pt3dadd(0, 0, 1015, 3)
}
// Connect section end points:
connect apic[0](0), soma[0](1)
// Set biophysical parameters:
forall {
 Ra = 100. // ohm * cm
 cm = 1. // uF/cm2
}
soma { insert hh }
apic {
 insert pas
 g_pas = 0.0002 // S/cm2
 e_pas = −65. // mV
}’’’)
~~~

We then proceed to set up the simulation of the model, by first importing different classes from LFPy and also other packages we may need:

~~~
**from** LFPy **import** Cell, Synapse, RecExtElectrode
**import** numpy as np
**import** matplotlib.pyplot as plt
~~~

First we instantiate the cell model, providing the name of the morphology file as well as simulation duration and resolution, and initial voltage for all compartments:

~~~
cell = Cell(morphology=morphology,
           tstop=100., # ms
           dt=0.05, # ms
           v_init=-65. # mV
          )
~~~

Next we instantiate the synapse with compartment index information, parameters for the chosen synapse mechanism, and set synaptic activation times:

~~~
synapse = Synapse(cell=cell,
                  idx=cell.get_idx(section=‘apic’)[-1],
                  syntype=‘ExpSyn’,
                  weight=0.05, # uS
                  tau=5., # ms
                  e=0., # mV
                  record_current=True)
synapse.set_spike_times(np.array([10., 15., 20., 25., 60.])) # ms
~~~

Then we instantiate the extracellular recording device, with a chosen value for the homogeneous extracellular conductivity and locations for the electrode contact points in space:

~~~
z = np.linspace(−100, 1100, 25) # um
electrode = RecExtElectrode(sigma=0.3, # S/m
                            x=np.zeros(z.size)+10,
                            y=np.zeros(z.size),
                            z=z)
~~~

Finally, we can simulate the model and predict the resulting extracellular potential:

~~~
cell.simulate(electrode=electrode)
~~~

The ‘simulate’ method of the cell does not return any data for storing, visualization etc., but sets various attributes of the instantiated class objects. For the present example, these are ‘cell.tvec’, ‘cell.somav’, ‘synapse.i’ and ‘electrode.LFP’. These are all numpy array types which can easily be plotted using matplotlib for example (see model output in Figure 1). The membrane depolarizations caused by the synapse current (*I*_syn_) result in action potentials at the soma (*V*_soma_). Both synaptic input and action potentials results in traveling features in the extracellular potential (image plot), towards the soma for synaptic input and from the soma for action potentials. These features are governed by the passive dendritic cable properties.

**Figure 1:**
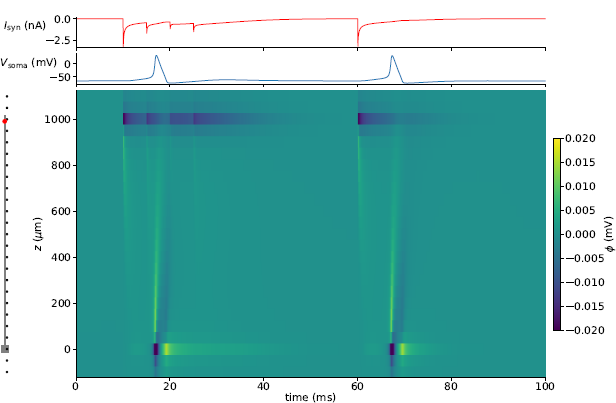
Single-cell example output. The morphology, synapse site (red dot) and recording locations (black dots) are shown on the left. The respective synapse current (*I*_syn_), soma potential (*V*_soma_) and extracellular potential (*ϕ*) are shown as function of time from top to bottom.

### 4.2 Electrostatic forward models

As the different electrostatic and magnetostatic forward models in LFPy are described in detail in Hagen et al. (2018), we here briefly summarize their main assumptions.

The relation between extracellular potentials and transmembrane currents are governed by volume conductor theory (Nunez and Srinivasan, 2006; Einevoll et al., 2013). At frequencies relevant for neuronal processes (below ten thousand hertz or so), the derivation of the volume conductor theory from first principles is simplified by application of the so-called quasistatic approximation to Maxwell’s equations (Hämäläinen et al., 1993, p. 426). The extracellular medium is in all cases assumed to be *ohmic*, which implies a linear and frequency-independent relation between currents and electric potentials (Pettersen et al., 2012; Einevoll et al., 2013; Miceli et al., 2017). Then, the simplest possible case is that of an *infinite homogeneous* (same in all locations) and *isotropic* (same in all directions) volume conductor (see e.g., Hagen et al. (2018, Sec. 2.2.1)). The medium can then be simply represented by a *scalar* extracellular conductivity, which assumes that local variations throughout the tissue from glia, dendrites, axons, cell bodies etc., are negligible when estimating the extracellular potential. This assumption of homogeneity appears valid for simulations mimicking recordings in for instance cortical gray matter (Goto et al., 2010).

For simple, analytically tractable circumstances, LFPy may also account for both inhomogeneous and anisotropic extracellular media. Inhomogeneities on the macroscopic level may arise at the interface between different tissue types such as white and gray matter, or between the gray matter and the conductive CSF on top of cortex. For such cases, LFPy can account for the effects of a discontinuous jump in conductivity by use of the so-called method of images (MoI), see Hagen et al. (2018, Sec. 2.2.2) for details. Another application of the MoI has been to mimic in vitro experimental setups, where jumps in conductivity occur between the petri dish, the tissue sample and saline layer covering the sample (Ness et al., 2015).

LFPy also supports forward-model predictions with homogeneous *and* anisotropic media, meaning that the electric conductivity may depend on direction but not location. This functionality may be utilized to mimic anisotropy in cortex (Goto et al., 2010), assuming that the typical orientation of dendritic and axonal fibers may affect the conductivity. For details on the implementation, see Hagen et al. (2018, Sec. 2.2.3).

In order to predict EEG signals, LFPy provides the analytically tractable 4-sphere head model (Nunez and Srinivasan, 2006; Næss et al., 2017). This simplified head representation assumes radial symmetry, and that the different electric conductivities of brain, CSF, skull and scalp are accounted for by corresponding concentric shells with homogeneous and isotropic conductivity. Neuronal sources assumed to be some distance away from the measurement site are represented by their *current dipole moment*, computed from the transmembrane currents in the neurons, see Hagen et al. (2018, Sec. 2.3.1).

The very same current dipole moment is utilized in order to compute distal MEG signals, computed using a special form of the corresponding magnetostatic Biot-Savart law (Nunez and Srinivasan (2006, Appendix C);Hagen et al. (2018, Eq. 16 in Sec. 2.3.5)). When computing magnetic fields in proximity to the neurons, the axial currents computed from the membrane voltage in each compartment are used instead of the current dipole moment. In both cases, negligible contributions from volume currents to magnetic signals are assumed, that is, axial currents within neurons are the main contributors to magnetic measures of brain activity (Hämäläinen et al., 1993).

### 4.3 Extensions to LFPy

As LFPy is written in Python, a transparent and easy-to-learn programming language, LFPy’s areas of application may with relative ease be extended to other measurement modalities or different forward models. Geometrically complicated variations in extracellular conductivity (including those introduced by the electrode device) can be accounted for using finite-element modeling (FEM (Logg et al., 2012; Lempka and McIntyre, 2013; Ness et al., 2015; Næss et al., 2017; Buccino et al., 2019)) in order to map transmembrane currents to extracellular signals. In a similar manner, current dipole moments computed using LFPy can be used with detailed head models constructed from MRI data (Bangera et al., 2010; DeMunck et al., 2012; Vorwerk et al., 2014; Huang et al., 2016). Frequency dependent conductive media may be accounted for by treating each Fourier component of recorded signals independently as pursued in Miceli et al. (2017).

One specific extension of the forward modeling scheme in LFPy is the so-called hybrid scheme for LFP predictions from point-neuron networks (Hagen et al., 2016). There, the simulation of ongoing spiking activity is first conducted offline using simplified neuron representations. The resulting spikes are used as activation times of synapses distributed onto geometrically detailed multicompartment neuron models, before each individual cell’s contribution to the LFP is simulated independently before summing up all contributions. Thus, the prediction of extracellular signals (LFP, EEG, MEG, etc.) of large neuronal networks is simplified by its disentanglement from simulations of recurrent network activity. Furthermore, the arduous effort of fine tuning parameters of recurrently connected networks of biophysically detailed neuron models is avoided.

Another extension to LFPy is detailed models of extracellular electric stimulation of neurons. LFPy do not include forward models for the effect of electrode stimulation currents onto neurons *per se*, but such can easily be constructed using the same assumptions as discussed above. These models will in fact be closely related to forward models mapping transmembrane current contributions to measured electrode potentials (through reciprocity). Given a prediction of the extracellular potential across the neuron’s outer surface, it can already be set as a cable-equation boundary condition using the `Cell.insert_v_ext` method, which relies internally on NEURON’s `extracellular` mechanism.

### 4.4 Deployability and documentation

LFPy is freely available (General Public License v3), runs on most common desktop computer operating systems (Linux, Unix, MacOS and Windows), and is presently tested with multiple versions of Python (2.7, 3.4-3.7). The software is made to run on computers ranging from laptops to HPC facilities. Its main dependencies upon installation are NEURON^2^ installed with bindings to Python (Hines et al., 2009), NumPy^3^, SciPy^4^, h5py^5^, mpi4py^6^ and Cython^7^.

### 4.5 Installation

NEURON can usually be installed using precompiled installers directly from its home-page (neuron.yale.edu), but certain systems (e.g., non-standard operating system configurations and HPC systems) may require compilation from source. More detailed instructions for installing both LFPy and NEURON on different common operating systems are provided through LFPy’s online documentation^8^.

The easiest way of installing stable releases of LFPy and its dependencies (except NEURON) is from the Python Package Index using the `pip` utility, by issuing in the terminal:

~~~
pip install LFPy --user
~~~

An existing installation can be upgraded by issuing:

~~~
pip install --upgrade --no-deps LFPy --user
~~~

Here, the `--no-deps` option disables attempts to update also other installed dependencies. The software can be removed by issuing:

~~~
pip uninstall LFPy
~~~

Alternatively, LFPy can be installed from source code hosted at GitHub.com/LFPy/LFPy. This repository contains the entire development history of LFPy using Git^9^. Installation of LFPy from source can be executed as follows:

~~~
**cd** <where to put LFPy sources>
git clone https://github.com/LFPy/LFPy.git
**cd** LFPy
pip install requirements.txt --user
python setup.py install --user
~~~

The GitHub homepage also offers the option to directly download compressed (zip or tar.gz) files with the source codes of LFPy.

In addition to the above requirements also matplotlib^10^ and the Jupyter Notebook^11^ are required to run all the example simulation scripts provided with LFPy. The different example files are provided with the LFPy source codes in the folder named `examples`.

### 4.6 Development and documentation

Active development and bug/issue tracking related to LFPy is conducted in public at GitHub.com/LFPy/LFPy. Fixes and new features are usually handled through Pull Requests, with automated build testing using Travis CI (travis-ci.org/LFPy/LFPy), and test coverage testing using Coveralls (coveralls.io/github/LFPy/LFPy). The full documentation for LFPy is provided at Read the Docs (lfpy.readthedocs.io).

## 5. Acknowledgements

The development of LFPy was supported by the European Union Horizon 2020 Research and Innovation Programme under Grant Agreement No. 720270 and No. 785907 [Human Brain Project (HBP) SGA1 and SGA2], the Norwegian Ministry of Education and Research through the SUURPh Programme and the Norwegian Research Council (NFR) through COBRA (grant # 250128), CINPLA and NOTUR -NN4661K.

Various individuals who have contributed to LFPy’s development are listed at GitHub.com/LFPy/LFPy/graphs/contributors.

## Footnotes

1 LFPy.readthedocs.io

2 neuron.yale.edu

3 numpy.org

4 scipy.org

5 h5py.org

6 bitbucket.org/mpi4py/mpi4py

7 cython.org

8 lfpy.readthedocs.io

9 https://git-scm.com

10 matplotlib.org

11 jupyter.org

